# MEDOC: A fast, scalable and mathematically exact algorithm for the site-specific prediction of the protonation degree in large disordered proteins

**DOI:** 10.1101/2024.10.08.617153

**Authors:** Martin J. Fossat

## Abstract

Intrinsically disordered regions are found in most eukaryotic proteins and are enriched in positively and negatively charged residues. While it is often convenient to assume these residues follow their model-compound pK_a_ values, recent work has shown that local charge effects (charge regulation) can upshift or downshift sidechain pK_a_ values with major consequences for molecular function. Despite this, charge regulation is rarely considered when investigating disordered regions. The number of potential charge microstates that can be populated through acid/base regulation of a given number of ionizable residues in a sequence, *N*, scales as ∼2^*N*^. This exponential scaling makes the assessment of the full charge landscape of most proteins computationally intractable. To address this problem, we developed MEDOC (Multisite Extent of Deprotonation Originating from Context) to determine the degree of protonation of a protein based on the local sequence context of each ionizable residue. We show that we can drastically reduce the number of parameters necessary to determine the full, analytic, Boltzmann partition function of the charge landscape at both global and site-specific levels. Our algorithm applies the structure of the *q*-canonical ensemble, combined with novel strategies to rapidly obtain the minimal set of parameters, thereby circumventing the combinatorial explosion of the number of charge microstates even for proteins containing a large number of ionizable amino acids. We apply MEDOC to several sequences, including a global analysis of the distribution of pK_a_ values across the entire DisProt database. Our results show differences in the distribution of predicted pK_a_ values for different amino acids, in agreement with NMR-measured distributions in proteins.

## Introduction

Electrostatic interactions are major driving factors for many molecular interactions^1–3^. In particular, ensembles of intrinsically disordered proteins (IDPs) are largely determined by the number and patterning of the charged residues in their sequences^4–8^. All charged amino acids, except for arginine^9–11^, can undergo acid/base charge regulation, by which the amino acid can lose or gain a proton from solution, thus altering its net charge^12–14^. The diversity of titratable amino acids, the number of possible charge microstates, and the conformations they populate make each titratable site potentially behave differently^15–18^. Measurements of both folded and disordered proteins suggest the effect of charge regulation can be significant^19–21^. An accurate model of electrostatic interactions for biomacromolecules requires an understanding of the underlying charge-state ensemble. Charge regulation can play a major role in the stabilization of protein structure^18,22,23^. However, our knowledge of both the impact of structure and local sequence context on the charge-state ensemble remains lacking, in large part because of the lack of theoretical tools to address the problem. A quantitative description of charge regulation and the sequence features that lead to it must be achieved to understand the relation between solution conditions, charge states, and protein structure and function.

Modeling the charge-state ensemble that can be populated by a polyacid is a complex problem due to the combinatorial explosion of possible charge microstates. A brute-force approach to this problem would involve first generating the list of all possible charge states, then evaluating the total free energy of each state by adding the contribution of each residue that releases a proton. In practice, the computational cost scales as *O*(2^*N*^), where *N* is the total number of ionizable residues in the sequence. A sequence of just 15 ionizable residues results in 32768 possible charge microstates, prohibiting a calculation of the full charge-state ensemble for larger systems. The computational cost is due both to the number of operations necessary to estimate the microstate weights and to the number of parameters, one for each microstate, which needs to be stored in memory.

To circumvent this problem, several approaches have been developed. One method is the use of machine learning to infer site-specific pK_a_ values of residues in a protein of interest^24–28^. However, pK_a_ predictors can hide much of the complexity that arises from electrostatic coupling. Such coupling can result in non-sigmoidal titration behavior, which these predictors cannot reproduce due to the reduction in the number of parameters to a single value per residue^29–31^. Another popular predictor for the charge-state behavior of disordered proteins, pepKalc^32^, circumvents this issue by calculating the full charge-state ensemble only for residues within a window, and for which the binary charge state of neighbors influence the protonation free energy. Residues outside the window are treated as fractional charges in a Gaussian chain model for their influence on the protonation free energy. This model allows for non-sigmoidal behavior by explicitly representing binary charge states within the window. However, it still relies on a simplification of the partition function to make the calculation tractable for large sequences.

In this paper, we describe our method for rapidly obtaining charge microstate weights from a database consisting of the free energy of ionization of each possible sequence context surrounding each ionizable residue type derived from simulations. We also show that we can compute the full set of parameters for data of both global and local scope for a protein exceeding 900 ionizable residues, without any combinatorial simplification. We first describe the method for rapidly obtaining the weights of charge microstates from a database consisting of the free energy of ionization of each possible context derived from the simulations. We show that the structure of the *q*-canonical ensemble can be leveraged not only to reduce the time to compute all possible weights, but also to compute the total protonation free energy associated with all of the states for both site specific and global protonation, without needing to compute individual weights. We are thus able to circumvent the combinatorial explosion because we use only the minimum number of parameters necessary to describe each type of experimental data.

## Materials and methods

The procedure within MEDOC uses a database of context dependent ionization free energies, where each of the ionizable amino acid deprotonation free energy contribution is modulated through neighbor amino acid type and neighbor sequence distance dependent weights that captures the modulation of ionization free energy by the sequence neighboring residues. These deprotonation free energies then used within an innovative algorithm to evaluate the contribution of all microstates to both the local and global scope observable that can be obtained from experiments. There are two main innovations that allows MEDOC speed and scalability. First, the structure of the *q*-canonical ensemble is used to wok with standard free energy that are independent of pH, but can be used to analytically recover the pH dependence, thus circumventing the need for an explicit discretization of the pH space. Second, rather than computing the free energy associated with each of the charge microstates, we instead compute the free energy associated with an ensemble of microstates that contribute to a particular observable. This allows to drastically reduced the number of mathematical operations necessary and reduce the number of individual free energies that must be kept in memory, thus allowing the prediction of large proteins while computing the analytically exact solution to the partition function effect on the observable.

Experimentally, two types of data can be used to estimate the charge-state ensemble, depending on whether their scope is global or local. Global measurements report on the average charge state of an entire protein, whereas local measurements offer information for specific residues. An example of a global measurement includes potentiometry titrations^33,34^, whereas local measurements include site-specific titrations using NMR experiments^20^. Importantly, neither measurement reports on the individual microstate weights, but rather reports on their average contribution to either the global or local protonation. For global measurement, the data report on the total degree of protonation of the chain. This can be written as:

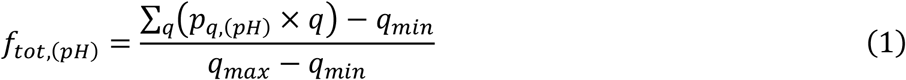

where *f*_*tot*,(*pH*)_ is the pH-dependent fraction of ionization, *q*_*min*_ and *q*_*max*_ are the minimum and maximum charges that the protein can populate, and *p*_*q*,(pH)_ is the pH-dependent fractional population of each charge mesostate that can be populated for a given net charge *q*. The population can be related to the canonical partition function:

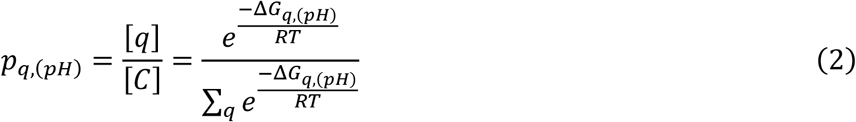

where [*C*] is the total concentration of the protein, and [*q*] is the total concentration of the protein in the mesostate of net charge *q*. In Eq. (2), *R* is the ideal gas constant, *T* is the temperature, and Δ*G*_*q*,(*pH*)_ is the pH-dependent free energy that can be obtained from the standard-state free energy 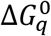 using the relation:

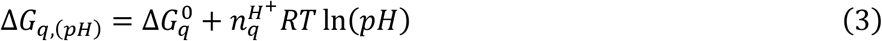

where 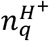 is the number of released protons for mesostate *q*. Note that we drop Δ from subsequent equations. In addition to simplifying the math, our notation is motivated by our choice of the reference state as the fully protonated state, such that *G*_ref_ ≡ 0 and Δ*G*_*μ*_ ≡ *G*_*μ*_ − *G*_ref_ = *G*_*μ*_. Similarly, unless specified by (pH) in the subscript, all free energies refer to the standard-state free energies. Thus, the number of parameters that such global measurements report on is *N* + 1, or one parameter for each mesostate, which is defined as the ensemble of charge microstates with a given number of bound protons.

In contrast, site-specific measurements report on the local degree of protonation, such that for each ionizable residue *r* represented in the data, we obtain a titration curve for the proton-bound state. We can define 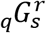 corresponding to the total free energy associated with states that have a particular residue *r* in a protonation state *s* for each mesostate *q*, enabling us to rewrite Eq. (2) as

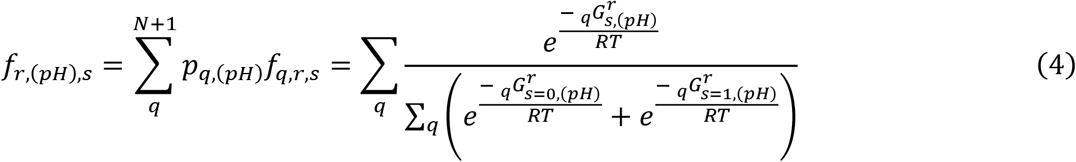

Which is the sum over all mesostates of the mesostate specific fraction of state *s* for a residue *r* (*f*_*q,r,s*_) times the pH-dependent probability of that mesostate (*p*_*q*,(*pH*)_). The full derivation of Eq. (4) is given in the supplementary information.

We denote the ionized (charged) state of each ionizable amino acid by its capitalized letter code and its non-ionized (uncharged) counterpart by its lower-case letter code. Thus, in the representative example in **Fig. 1**, “e” (protonated glutamic acid) carries a net charge of 0, and “E” (deprotonated glutamic acid) carries a net charge of -1. The context-dependent ionization free energy of each ionizable residue in the sequence can be obtained simply by summing the different context-specific ionization free energies. The standard free energy of ionization of a glutamic acid dipeptide, where both amino acids carry a net charge of -1, denoted by EE, is therefore the sum of the free-energy change associated with the transitions ee⟶Ee and Ee⟶EE, following a thermodynamic cycle.

**Figure 1:**
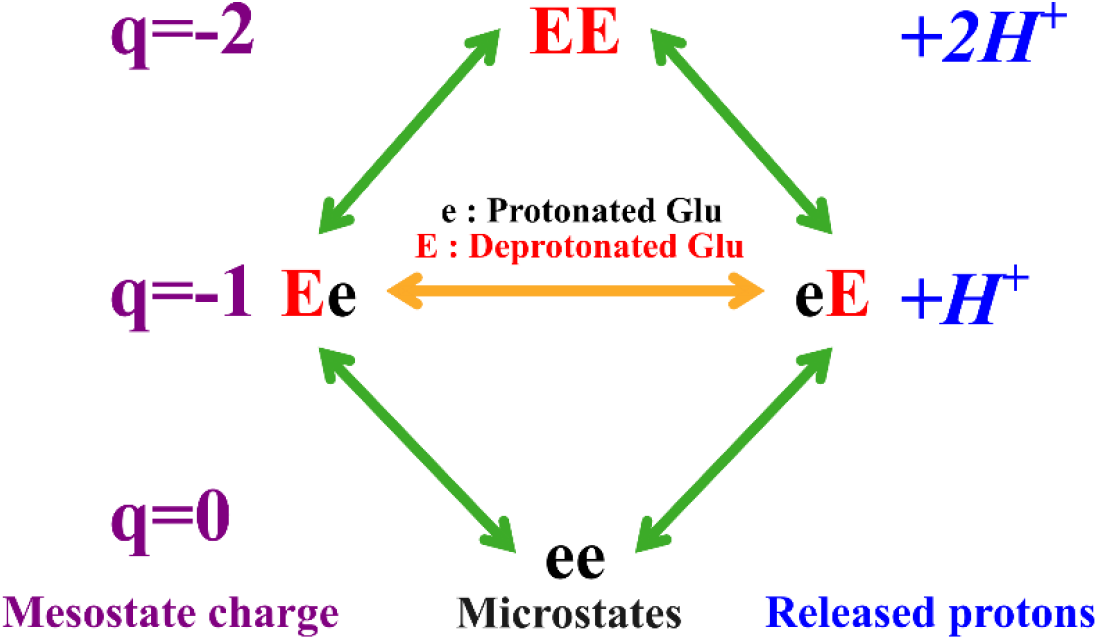
Overview of the q-canonical ensemble for a small peptide of two glutamic acids. The mesostate charges are indicated in purple, the number of released protons for each mesostate is indicated in blue.

This cycle is summarized in **Fig. 1**, where we show the q-canonical clustering of charge microstates into mesostates.

Our approach rests on a pre-generated database of context-specific free energies of ionization that we use to estimate the free energy of ionization of each ionizable residue. To do so, we define a window as the number of neighbors on each side of the ionizable amino acid of interest. The effect of different window sizes is discussed below. For each residue in each sequence, this context-specific ionization free energy is estimated using Hamiltonian exchange free energy *q*-canonical ABSINTH^35,36^ simulations. The assumption that residues outside of the window do not significantly contribute to the free energy of ionization implies a lack of sequence-distance interactions.

For each microstate the same free energy can be obtained using different thermodynamic paths, which correspond to the different orders in which the amino acid can lose its proton, as shown in **Fig S1**. We choose always to employ the thermodynamic path in which the contribution to the microstate deprotonation free energy of each ionizable residue is added from the N-terminal to the C-terminal residue in the sequence. Choosing this thermodynamic path limits the number of context-specific deprotonation free energy simulations necessary for the full description of the charge ensemble, because we do not need to simulate any of the contexts in which a right-side neighbor has lost its proton. The individual context-dependent free energy of ionization can be obtained in two ways: either by direct simulation of the peptide excised from the sequence, which we call here the *explicit context database*, or by estimation using sequence heuristics, which we refer to as the *implicit context database*. In the explicit context database, we perform free energy calculations directly on the sequence context of interest. For example, obtaining the free energies of deprotonation for microstates of a sequence of 5 Glu requires free energy simulations of each of the possible contexts for every amino acid in the microstate that has released a proton, as depicted in **Fig. 2A**. In the implicit method, we estimate the individual contribution of each pattern to the weight of a microstate and rely on a database of free energy estimates based on the assumption of additivity between the contribution of each neighbor to the ionization free energy of the central residue in the pattern. A schematic of this process can be found in **Fig 2B**. A comparison between the explicit and implicit methods resulting context dependent deprotonation free energy, along with results from a q-canonical ABSINTH simulation of the full peptide, is shown in **Fig. 2C**. Full details are available in the supplementary information.

**Figure 2:**
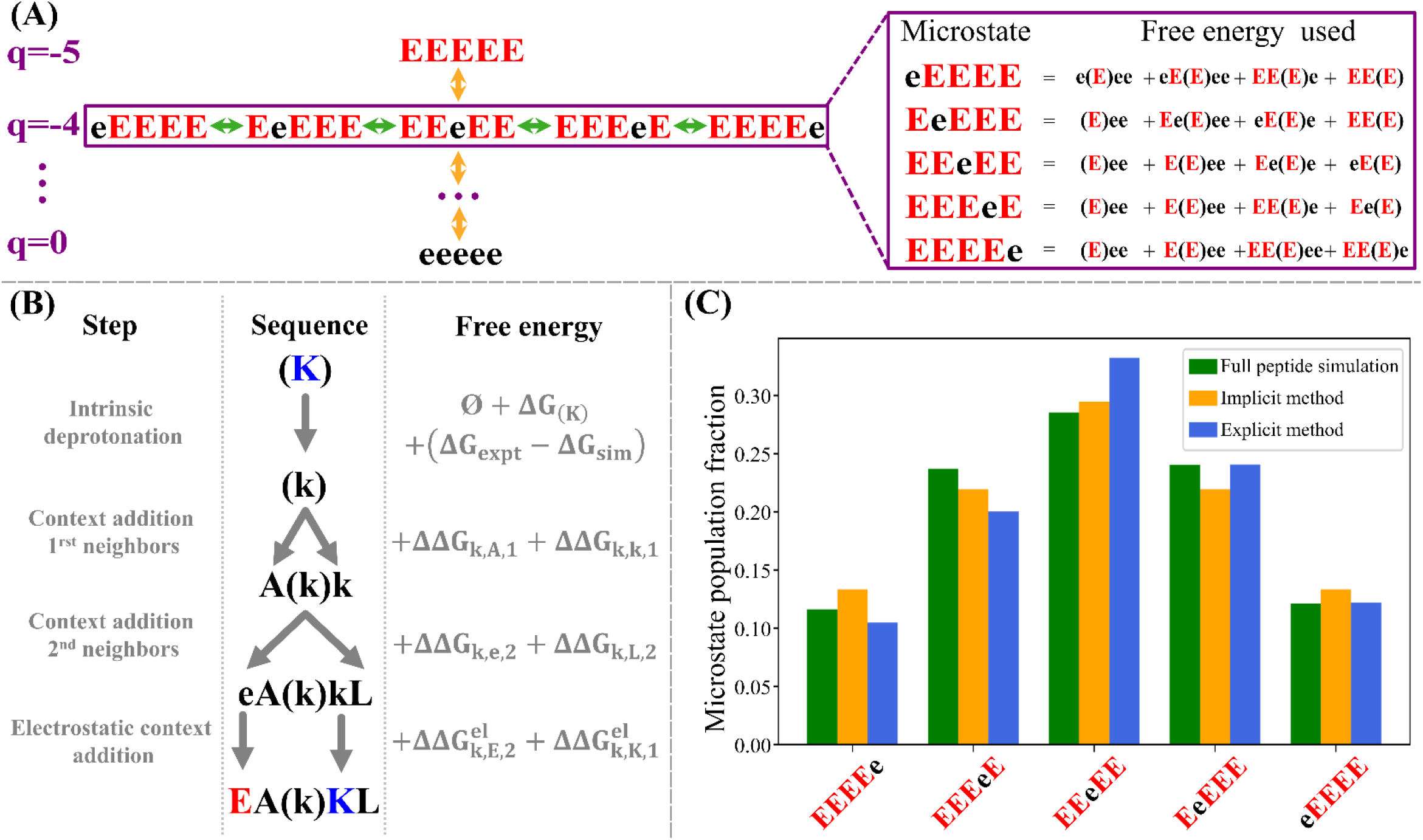
(A) Representation of the mesostate organization of the q-canonical ensemble (left) and schematic of the context-dependent ionization free energy for a window size of 2 as described in the text (right). The amino acid in parentheses is the amino acid for which the deprotonation free energy is added to the microstate free energy. The lower-case letters represent the protonated state of Glu. The ionization free energy for each context can be computed by either the explicit or implicit methods as described in the text. (B) Schematic of the implicit method for obtaining the deprotonation free energy of individual contexts. (C) Bar plot showing the intra-mesostate populations derived from the full simulation (green), the implicit database method (yellow), and the explicit database method (blue).

MEDOC uses two algorithmic procedures in combination to reduce the number of necessary calculations for obtaining the full set of parameters to extract the global and local protonation curves. The first procedure, dynamic building of the microstate ensemble, is used to remove the redundant operations when computing microstates that share the same protonation state up to the residue that is currently being evaluated. The second procedure, determination of the ensemble weights, builds on the first and is used to only keep in memory the global parameters that will be necessary to compute the full partition function at the end of the algorithm, without storing the contribution of individual microstates. Using these procedures in combination allows us to drastically reduce both the number of operations and the number of parameters needed to store in memory, which are limiting factors in the full determination of the protonation partition function.

We can reduce the number of steps necessary for the determination of the free energy of each microstate, by considering sequence redundancy in the computation. For example, the sequences Eee and EeE both have the same free energy associated with the protonation states of the first two residues. This means we can compute the free energy of the microstates as a sum of the same free energy associated with an Ee context, plus the free energy associated with the residue following the context. If we repeat this procedure, we can obtain each of the microstate weights without computing the same context twice. We illustrate this idea in **Fig. 3**. Removing these redundant calculations leads to improved efficiency for the algorithm. However, the number of microstate weights that are necessary to save in memory still limits applications that are based solely on this method. The procedure presented in the following paragraph solves this issue.

**Figure 3:**
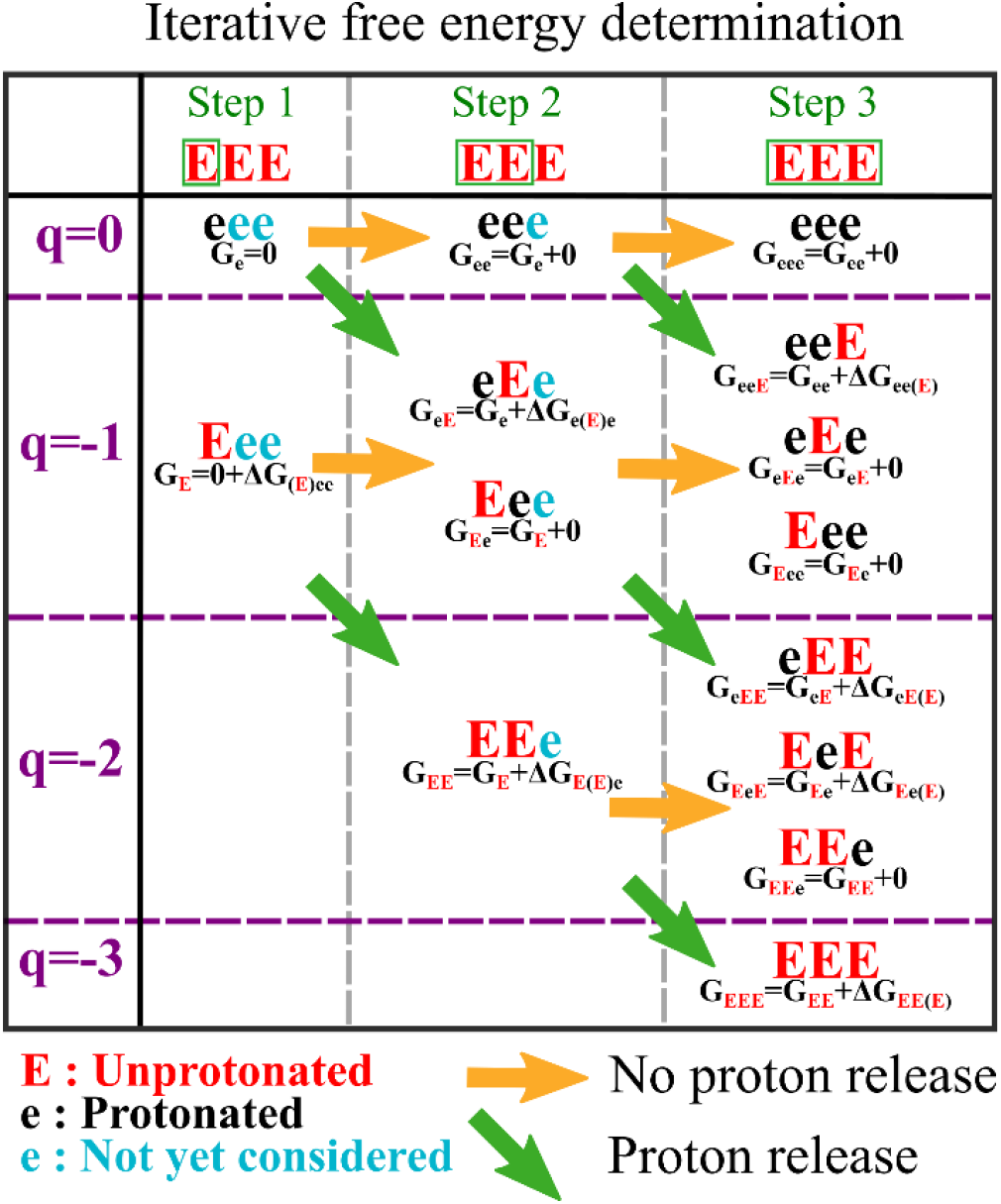
Depiction of the dynamic building procedure to efficiently generate charge microstate standard free energy.

To improve the capability of the algorithm to analyze sequences with a large number of ionizable amino acids, we developed a method for reducing the number of mathematical operations upon the addition of context-dependent protonation free energies. This allows us to store the contribution of all microstates to the partition function, without storing the free energy of individual microstates. To make the notation clearer, we define the overall protonation free energy corresponding to *n* microstates:

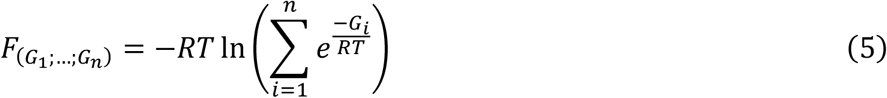

We define the contribution of the overall free energy to the context for each mesostate *q* at step *t* during the iterative procedure, which we denote 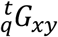, where *x* and *y* are 0 or 1, corresponding to the protonation state of the last two ionizable residues accounted for so far. If we define *N*_*xy*_ as the number of microstates that end in a given context, we obtain:

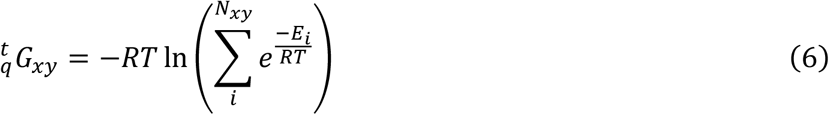

We then propagate this free energy by considering the next residue in the sequence in cases where the residue either releases or does not release a proton. If a proton is released, the new energy of each microstate is *E*_*i*_ + Δ*E*_*xy*_, such that:

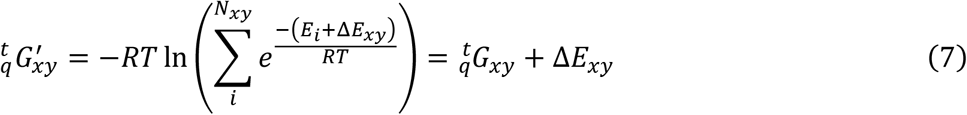

If the newly considered residue does not lose a proton, no deprotonation free energy is added, and 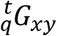 does not change. Thus, for each step, we propagate the context-specific free energy by recombining the contribution of the contexts in the previous step, using the following equations:

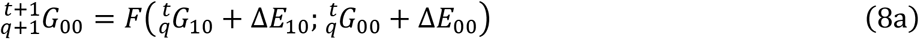

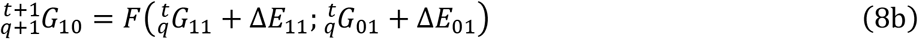

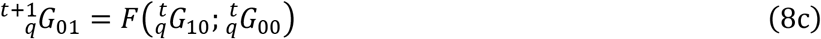

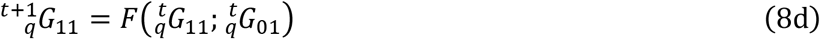

When this procedure is applied to all ionizable residues in the sequence, we recover the overall free energy associated with all microstates of a given mesostate *q* using:

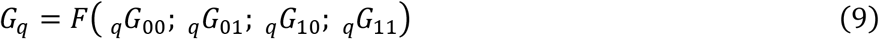

Using Eq. (9), we compute the total contribution to the overall degree of protonation of the protein defined in Eq. (1) for all microstates while accounting for the specific context-dependent deprotonation free energy, without having to explicitly represent the microstates energies themselves. The overall algorithm is depicted in **Fig. S2** and **Movie S1** for two contexts for a single step. A pseudo code version of the algorithm is depicted in **Fig. 4**.

**Figure 4:**
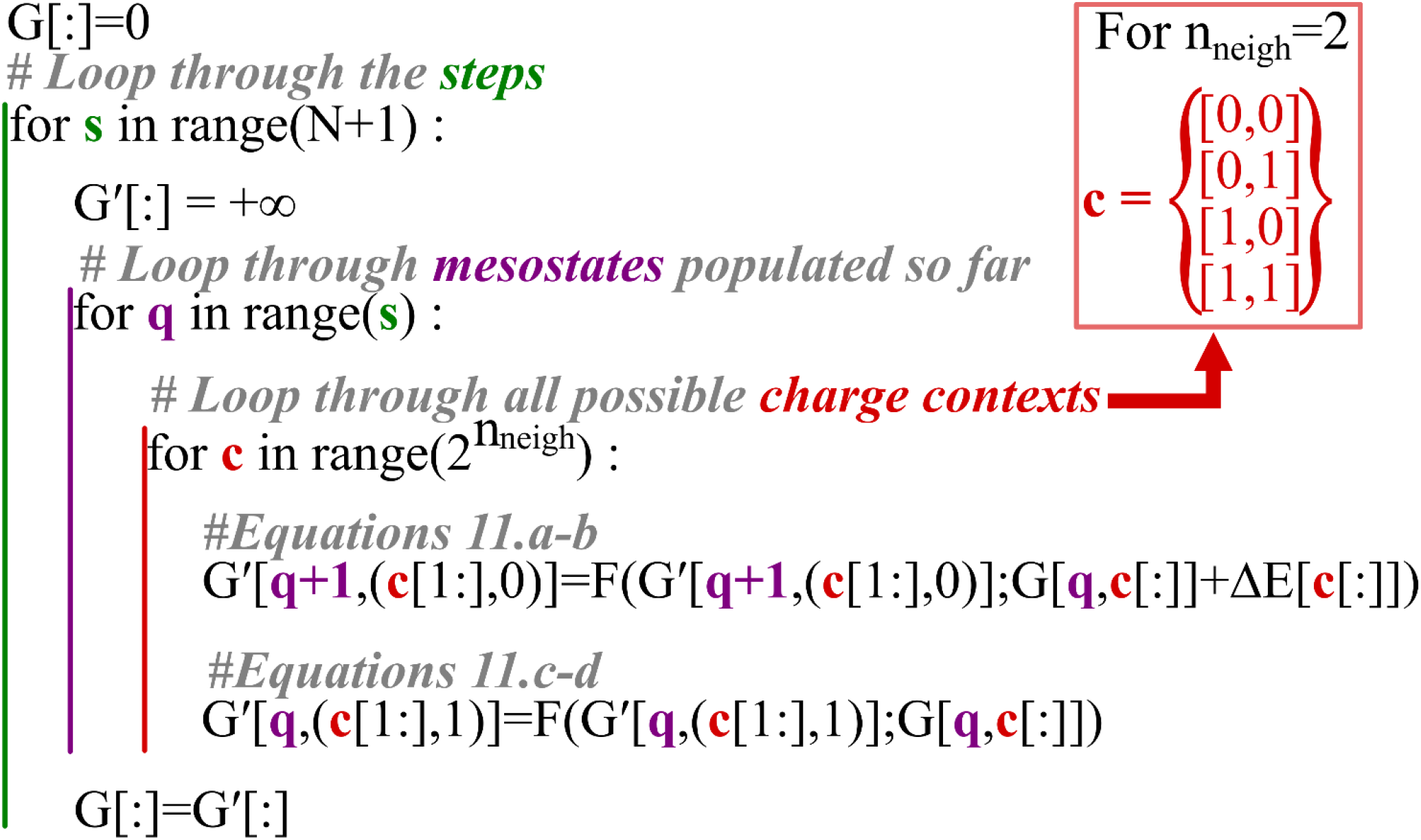
Pseudo code depicting the ensemble weight determination method. The algorithm loops through N+1 steps (green), looping through all mesostates that are populated so far as depicted in **Fig. 3** (purple). The algorithm then loops through the possible charge context, which are the charge states if the two last residues. The indices of G and G’ are the charge mesostates and the charge context. (c[1:],0) indicates the indices of the last n_neigh_- 1of the context and a 0 are the indices of G[q]. A more detailed description of the processes is available in **Fig. S2**.

To make site-specific predictions, we compute a residue-specific partition function for each ionizable residue in both their proton bound and unbound states, i.e., a 1 or a 0. We note the free energy of residue position *p* in the charge state *s* after *t* steps of the algorithm for charge mesostate *q* as 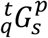. This quantity is itself sub- divided into four quantities that represent the protonation states of the last two residues that have been considered. Mathematically, this can be expressed as:

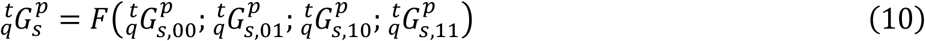

This decomposition allows us to keep in memory all possible context-specific contributions to each of the residues and each layer in each state. For each step of the algorithm, we compute the free energy of the new combination of states from the previous one for each of the possible contexts for each position:

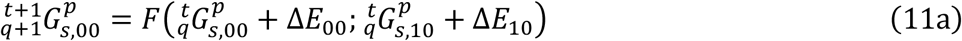

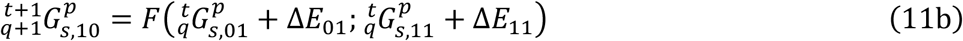

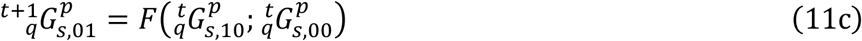

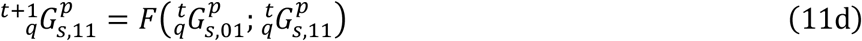

At each step, a new ionizable residue is considered, and we need to inherit the free energy from the previous residue:

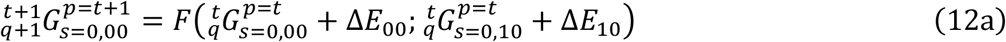

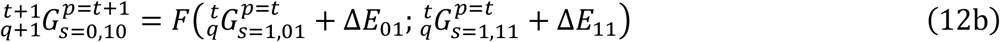

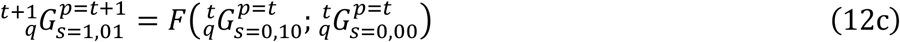

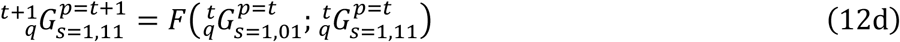

If this procedure is repeated until *t* = *N* + 1, Eq. (10) corresponds to the standard free energy of all microstates that have any given residue in both the protonated and unprotonated states for each mesostate, which can be used to compute the residue specific profile using the partition function defined in Eq. (4).

The input parameters for our prediction method were obtained using the ABSINTH implicit solvation model^35^ as implemented in the CAMPARI simulation engine. As these simulations needed to be independent of any specific secondary structure, we performed all simulations at a temperature of 400K, thereby flattening the conformational landscape. Each simulation was set up as a Hamiltonian replica exchange procedure interpolating between the fully protonated and fully deprotonated state for the central residue of the pattern, as described previously^36^. The free energies were then extracted using the multistate Bennett acceptance ratio (mBAR)^37^ procedure as implemented in the pymbar^38^ Python library on the output cross free energy for all replicas. We note that although the original simulations were performed at 400K, the mBAR procedure works directly with input energies, which are independent of temperature. Given that we did not use any temperature-dependent force field parameters in our simulation, the temperature only alters the acceptance ratio, making all conformations more likely.

The implicit free energy of deprotonation parameters were rescaled in order to improve the agreement with published site specific pK_a_ values (**Fig. 5C**). This rescaling was done by separately adding a factor to the electrostatic and intrinsic free energy of deprotonation contribution separately for each of the ionizable amino acids. Details on how this was done are available in the supplementary.

**Figure 5:**
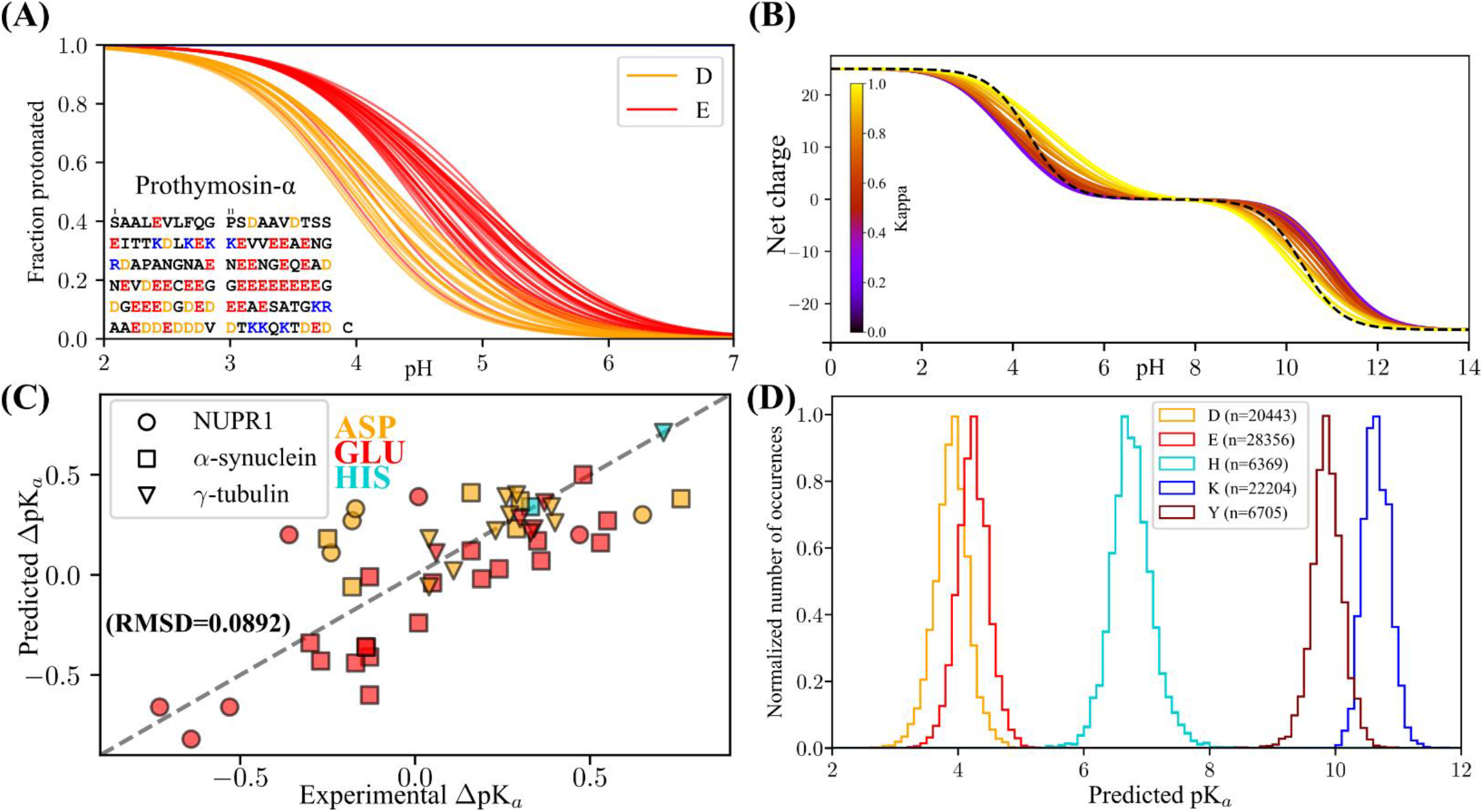
(A) Site-specific prediction of the fraction protonated for each the glutamic and aspartic acid residues in Prothymosin-α as a function of pH. (B): Net charge vs pH as predicted for all thirty of the fixed composition (25 Glu and 25 Lys) Kappa sequence variants presented in Das et al. 2013^4^. The Kappa value represents the degree of charge segregation, such that the sequence with a maximum Kappa of 1 is E_25_K_25_, and the sequence with the minimum Kappa is (EK)_25_. The dotted black line corresponds to the curves assumed from canonical unshifted pK_a_ values from Platzer et al.^40^. (C) Predicted vs Experimental pK_a_ for 3 proteins for which pK_a_ values have been measured by NMR^16,41,42^. The label used is protein specific, which the color is residue type specific, as indicated in the figure. (D) Distribution of predicted pK_a_ values using MEDOC for 5925 sequences (84077 individual ionizable sites) from the DisProt database^43,44^. The number of values for each amino acid is written in the legend.

**Figure 6:**
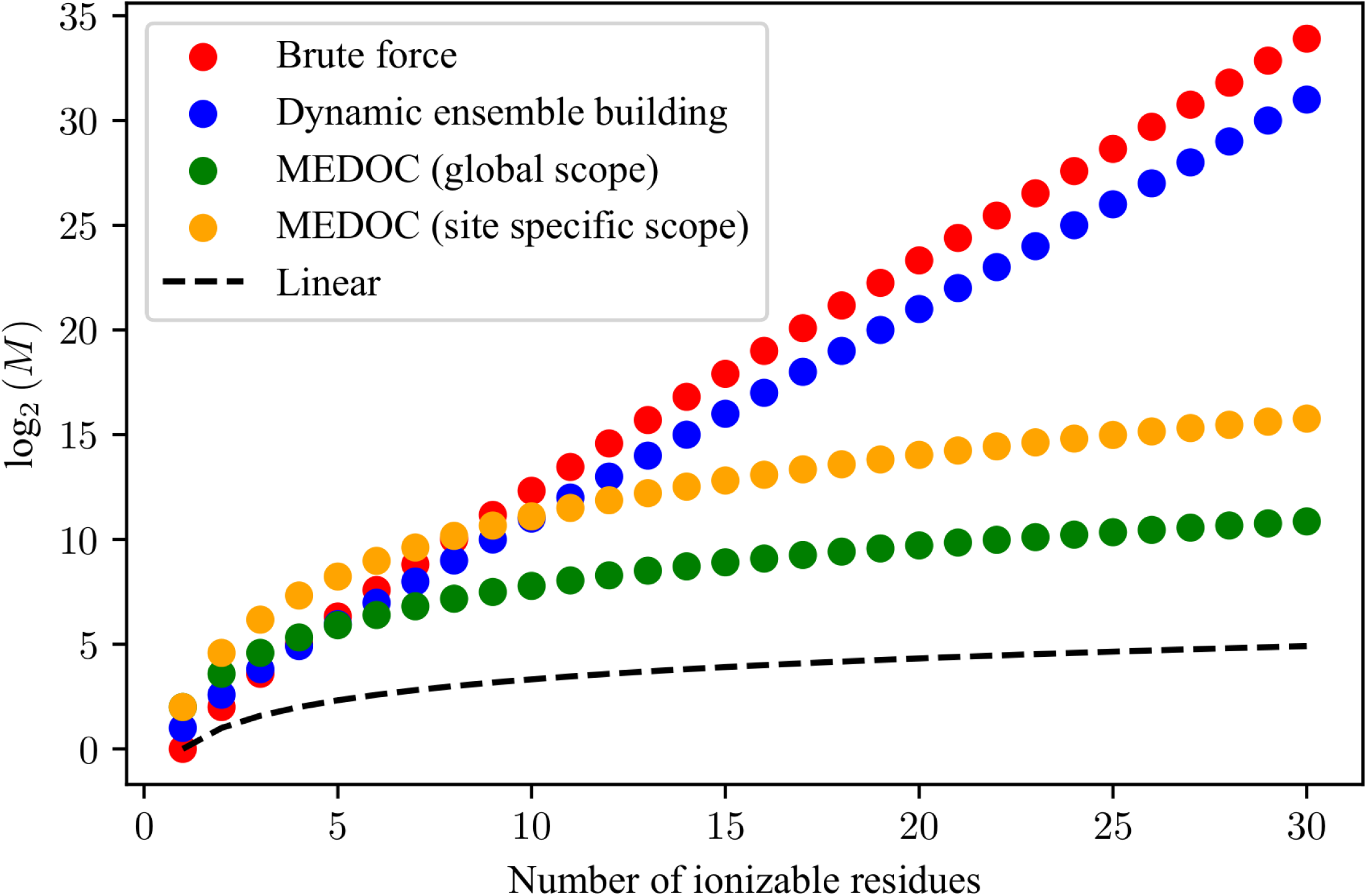
Plot of the log_2_ of the total number of mathematical operations necessary (M) for the determination of the full partition function as a function of the number of ionizable residues for a window size of 4. The calculations behind this graph are detailed in the supplement.

## Results and discussion

We presented two methods to obtain the context-specific free energies of ionization that serve as starting values for the prediction of the full partition function; the context database method and the additive free energy method. Each method has advantage and disadvantages. The context database method makes use of a database of context-specific free energy of ionization. While this is more accurate, the number of patterns for which we need to obtain a free energy is extremely large, as discussed in the supplementary information. For this reason, we present it here only for comparison with the additive method. By contrast, in the additive free energy method, any context can be deducted from 900 simulations for a window size of 6. We show that the two methods are in good agreement with one another by comparing the microstate free energy within a mesostate in **Fig. 2C**. Due to the major advantages of the implicit method, this work reports results computed using this approach.

To examine the effect of the window size used to calculate local charge effects, we computed the apparent net charge for an (E)_25_(K)_25_ peptide as a function of pH over several different window sizes (**Fig. S3**). This revealed that beyond a window size of 3, increasing the window size had a vanishingly small impact on the predicted charge state, with the curves assymptoting to a window size of 5. Consequently, we chose a window size of 5 moving forward. Additional window size tests are available in the supplement.

We first investigated predicted pK_a_ effects on the strong polyelectrolyte Prothymosin α (ProTα), a well-studied negatively charged polyelectrolyte involved in histone chaperoning^39^. We examined the predicted pK_a_ values for each individual negative titratable residue in ProTα (**Fig. 5A**). We next examined applied MEDOC to investigate the charge state prediction for the commonly used Kappa (κ) variants^4,7^. These sequences were designed to showcase the dependence of conformation on the distribution of oppositely charged amino acids. Each of the 30 variants is composed of 25 Lysines and 25 Glutamic acids, ranging from a sequence of alternating charges, (EK)_25_ (κ=0), to two stretches of opposite charges, (E)_25_(K)_25_ (κ=1). The conformation of these sequences go from an expanded globule for the (EK)_25_ variant to more compact hairpin like structures for the (E)_25_(K)_25_ variant^4^. Because the importance of neighboring charges on the determination of ionization free energy, the same factors contributing to the difference in conformation also affect the charge state of the variants. Our prediction shows that the segregation of like-charges results in an increase in the charge state heterogeneity around neutral pH, and a decrease in proton binding cooperativity, which can be seen in the decrease in the slopes of the curves as a function of Kappa (**Fig. 5B**).

We have plotted the predicted vs experimental pK_a_ values obtained by NMR^16,41,42^ in (**Fig. 5C)**. In this plot RMSD=0.0892 indicating a strong agreement of the prediction relative to the experiment, despite the absence of consideration for conformation in our algorithm.

The throughput of MEDOC allows for its deployment of our methods on large sequence databases. We have deployed our method to all proteins constituting the DisProt database^43,44^, for which we have extracted the pK_a_ values for each of the 84077 ionizable residues from the 5925 sequences in the database, and plotted their distribution by residue type in **Fig. 5D**.

A central feature of MEDOC is the ability to compute full charge distributions for disordered proteins large numbers of titratable residues. To motivate this, we examined the computational gains for our algorithm by calculating the number of necessary mathematical operations to obtain the full partition function using the full calculation without elimination of redundant microstates (red), the full calculation using only the dynamic building of the microstate ensemble procedure (blue), and the full MEDOC procedure for the site-specific (yellow) and global (green) cases. Details on the calculations of the number of necessary mathematical operations are available in the supplementary information.

The limitations of MEDOC are twofold: first, in its current version, the context parameters used as an input are derived from free energy calculation in the implicit solvent force field ABSINTH. Accordingly, the derived free energies may not be accurate, given the use of the additivity paradigm in this and other implicit solvent force fields. Second, the assumption that residues outside the window do not contribute to the deprotonation free energy is only realistic for highly expanded sequences. In that sense, MEDOC, as its name implies, is an estimation of the sequence context contribution to the extent of deprotonation in the absence of the influence of conformation. Despite these limitations, MEDOC can be used to obtain a better prior of the charge ensemble of a protein, and is capable of tackling large sequences (N>900) with a speed that allows for predictions of proteome wide datasets. We hope to address these limitations in future work.

## Conclusion

Simulations of biomacromolecules in large part rely on the fixed charged assumption that only one state dominates the charge-state ensemble at any given pH value. While historically this assumption has been made purely for practical reasons, recent work has highlighted the importance of charge regulation and its impact on structure^45^. Our method, MEDOC, circumvents the large number of possible charge microstates when solving for the full partition function, by nonetheless considering all the microstate weights contributions, thereby making the problem of assessing the full charge landscape of proteins tractable even for large sequences. In contrast to existing methods, MEDOC proposes to determine the full partition function with an analytically exact solution, while having a computational cost scaling as ∼*N* for the global scope and ∼N^2^ for the site-specific scope instead of ∼2^*N*^. This is achieved by a combination of algorithmic methods and the use of both a protonation free-energy database and the minimum number of parameters necessary for a full determination of the data reported by experiments. We hope that MEDOC will help the community to explore the potential effect of acid/base charge regulation. This is of particular relevance in the light of recent measurements of significant differences in pH values across the different phases of the nucleolus^46^. Furthermore, the proteins that constitute this nuclear body show a particular bias towards a high fraction of charged residue and high charge segregation (Kappa). As we showed, these sequences are more likely to undergo acid/base charge regulation, which could indicate an evolutionary selection for sequences prone to acid/base charge regulation. We will use MEDOC to sample the sequence space of these proteins and identify sequence features that are likely to result in large acid-base charge regulation effects. This will help the community at large by providing new insight into charge states and pH dependence of proteins.

Data availability and distribution:

MEDOC is available on GitHub (https://github.com/martinfossat/MEDOC_public.git).

## Acknowledgement

This work was supported by Air Force Office of Scientific Research under Grant No. [FA9550-20-1-0241], US National Institutes of Health under Grant No. [R01NS121114]. We also acknowledge additional support from Rohit Pappu in conduction this work, as well as Samuel Cohen for his advice on redacting the manuscript.

