## Supplement for "MEDOC: A fast, scalable and mathematically exact algorithm for the site-specific prediction of the protonation degree in large disordered proteins"

Martin Fossat

**Combining free energies**

We can divide these contributions into mesostate-specific free energies. Assuming we have two distinct groups of microstates of populations $p_1$ and $p_2$, with respective free energies $G_1$ and $G_2$, then:

$$p_{1+2} = p_1 + p_2 \tag{S1}$$

$$\frac{W_{1+2}}{W_{tot}} = \frac{W_1}{W_{tot}} + \frac{W_2}{W_{tot}} \tag{S2}$$

Where $W_i$ is the Boltzmann weight of microstate $i$. Given that we can assign a free energy to a state that we define as the combination of state 1 and 2, which has the free energy:

$$G_{1+2} = -RT \ln\left(e^{\frac{-G_1}{RT}} + e^{\frac{-G_2}{RT}}\right) \tag{S3}$$

Which is how we derive equation 5 in the main text

**Site specific measurements equation derivation**

As mentioned in the main text, site specific measurements report on the local degree of protonation, such that for each ionizable residue $r$ represented in the data we obtain a titration curve for the protonated fraction. We define $S_{\mu,r}$ as protonation state of residue $r$ in microstate $\mu$ such that $S_{\mu,r} = 1$ if the proton is bound, and $S_{\mu,r} = 0$ if not. The residue specific fraction protonated can thus be expressed as:

$$f_{r,(pH),S=1} = \sum_{\mu}^{2^N} S_{\mu,r} \frac{[\mu]}{[C]} \tag{S4}$$

Or, for the unprotonated fraction:

$$f_{r,(pH),S=0} = \sum_{\mu}^{2^N} (1 - S_{\mu,r}) \frac{[\mu]}{[C]} \tag{S5}$$

We can recast this in the $q$-canonical formalism, by grouping the microstates into constant net charge mesostates. In this case, the number of microstates within each mesostate is $N_{\mu,q} = \binom{N}{n_q^{H^+}}$ with $n_q^{H^+}$ being the number of protons bound to the chain in that mesostate, such that:

$$\sum_{q} \sum_{\mu}^{N_{\mu,q}} 1 = 2^N \tag{S6}$$

$$f_{r,(pH),S=0} = \sum_{q} \frac{[q]}{[C]} \sum_{\mu}^{N_{\mu,q}} (1 - S_{\mu,r}) \frac{[\mu]}{[q]} \tag{S7}$$

Which we can write as:

$$f_{r,(pH),s} = \sum_{q}^{N+1} p_{q,(pH)} f_{q,r,s} \qquad (S8)$$

With:

$$\mathrm{f}_{\mathrm{q,r,s=0}} = \sum_{\mu}^{\mathrm{N}_{\mu,\mathrm{q}}} \left(1 - \mathrm{S}_{\mu,\mathrm{r}}\right) \frac{[\mu]}{[\mathrm{q}]} \qquad (S9)$$

And

$$f_{q,r,s=1} = \sum_{\mu}^{N_{\mu,q}} S_{\mu,r} \frac{[\mu]}{[q]} \qquad (S10)$$

The corresponding free energy each state within a mesostate *q* with residue *r* in state *s* is:

$${}_{q}G_{s}^{r} = -RTln\left(f_{q,r,s}\right) \qquad (S11)$$

We can obtain the pH dependent free energy using equation 3. Because the sum of all microstates in a mesostate that have a residue in state 1 and 0 is all the states in that mesostate, we can write:

$$f_{q,r,s} = \frac{e^{\frac{-{}_{q}G_{s,(pH)}^{r}}{RT}}}{e^{\frac{-{}_{q}G_{s=1,(pH)}^{r}}{RT}} + e^{\frac{-{}_{q}G_{s=0,(pH)}^{r}}{RT}}} \qquad (S12)$$

We note that this fraction can also be expressed using the pH independent ${}_{q}G_{s}^{r}$ because the $n_{q}^{H^{+}} G^{H^{+}}$ simplifies due to all states within that mesostates having released the same number of protons, however, the full notation is important for following equations. We can also rewrite equation 2:

$$\mathrm{p}_{\mathrm{q,(pH)}} = \frac{[\mathrm{q}]}{[\mathrm{C}]} = \frac{\mathrm{e}^{\frac{-{}_{\mathrm{q}}\mathrm{G}_{\mathrm{s=1,(pH)}}^{\mathrm{r}}}{\mathrm{RT}}} + \mathrm{e}^{\frac{-{}_{\mathrm{q}}\mathrm{G}_{\mathrm{s=0,(pH)}}^{\mathrm{r}}}{\mathrm{RT}}}}{\sum_{\mathrm{q}} \mathrm{e}^{\frac{-{}_{\mathrm{q}}\mathrm{G}_{\mathrm{s=1,(pH)}}^{\mathrm{r}}}{\mathrm{RT}}} + \mathrm{e}^{\frac{-{}_{\mathrm{q}}\mathrm{G}_{\mathrm{s=0,(pH)}}^{\mathrm{r}}}{\mathrm{RT}}}} \qquad (S13)$$

Which means equation S8 can now be expressed as:

$$f_{r,(pH),s} = \sum_{q} \left( \frac{e^{\frac{-{}_{q}G_{s=1,(pH)}^{r}}{RT}} + e^{\frac{-{}_{q}G_{s=0,(pH)}^{r}}{RT}}}{\sum_{q} e^{\frac{-{}_{q}G_{s=1,(pH)}^{r}}{RT}} + e^{\frac{-{}_{q}G_{s=0,(pH)}^{r}}{RT}}} \times \frac{e^{\frac{-{}_{q}G_{s,(pH)}^{r}}{RT}}}{e^{\frac{-{}_{q}G_{s=1,(pH)}^{r}}{RT}} + e^{\frac{-{}_{q}G_{s=0,(pH)}^{r}}{RT}}} \right) \qquad (S14)$$

And thus :

$$f_{r,(pH),s} = \sum_{q} \frac{e^{\frac{-{}_qG^r_{s,(pH)}}{RT}}}{\sum_q e^{\frac{-{}_qG^r_{s=1,(pH)}}{RT}} + e^{\frac{-{}_qG^r_{s=0,(pH)}}{RT}}} \quad (S15)$$

Which is equation 4 in the main text. Thus, by obtaining the free energies ${}_qG^r_s$ for every combination of $q$, $r$ and $s$, which is the goal of our method, allows us to represent the full pH titration of each site with a minimum number of parameters with no simplification to the expression of the partition function.

### Effect of window size on results

In addition to **Fig. S6** mentioned in the main text, we looked at the context dependent ionization free energy parameters used for the additive context-based prediction (**Fig. S4** and **Fig. S5**). We can see that even at greater window lengths, the parameters may still not be zero, in particular for the electrostatic contribution. Furthermore, the electrostatic parameters of the 7$^{th}$ neighbors show that at this sequence length, which approaches typical protein blob size[1], the magnitude of the influence starts increasing due to the chain looping back on itself. Because we explicitly do not want sequence specific conformations to influence the parameters, we chose a window size of 5 in this work, unless specified otherwise. Because our model propagates the contribution to the microstate energies, distant residues can influence the protonation behavior if they are "connected" though a network of ionizable residues with overlapping contexts. This is because these residues may not influence each other's deprotonation free energy directly, but still affect the microstate free energies and thus the partition function. We illustrate this by showing the difference in the residue specific titration curve for a residue for two sequences that only differ by a single residue outside window size of 2 in **Fig. S6**.

### Explicit method database size

The explicit method for obtaining context specific deprotonation free energies is limited by the size of the sequence space. The total number of sequence contexts that can be obtained is expressed as :

$$\mathrm{S_z} = \mathrm{n_1^{n_{neigh}} n_2 n_3^{n_{neigh}}} \quad (S16)$$

If we consider Asp, Glu, His, Tyr and Lys as ionizable, and taking into account that the protonation states of the two rightmost neighbors are fixed by choice of the thermodynamic path, then $n_1 = 26$, and $n_3 = 20$, $n_2 = 5$. Thus, the database size is $S_z = 1352000$ for considering only two neighbors on each side. We can look at the Disport database to quantify the number of pentapeptides that would be necessary for a full description of the database. This is done by identifying each of the ionizable amino acids and its pentapeptide context, and then determining the number of possible contexts for residues that have ionizable left-side neighbors. We find that there are 26537 ionizable residues at unique positions in the Disprot database. Because of the redundancy in pentapeptide patterns that surround these residues, there are 20838 unique pentapeptides. If we account for the number of possible charge contexts, this number goes up to 35555. This is however still a very large number of free energy calculation, which is why we choose to use the implicit method in most of the manuscript and why the explicit method cannot be released to the public.

### Implicit context free energy determination

To estimate the context specific deprotonation free energy using simple heuristics, we use a small database of symmetrical peptides with an increasing number of identical neighbors that we denote $Z_n(X)Z_n$, where X is the ionizable amino acid for which we obtain the ionization free energy, and z can be any of the 20 common amino acids. We then used this database to compute the influence of the ionization free energy on each position for each amino acid/ionizable amino acid pair. The influence of the addition of an amino acid at a position sequence distance of *n* is defined as the difference between the ionization free energy of $Z_n(X)Z_n$ and $Zz_{n-1}(X)z_{n-1}Z$ such that:

$$\Delta\Delta G_{X,Z,n} = \frac{\Delta G_{z_n(X)z_n} - \Delta G_{z_{n-1}(X)z_{n-1}}}{2} \qquad (S17)$$

For the ionizable residues, a separate term is used to represent the change in ionization free energy upon ionization of a neighboring residue. This term is obtained by calculating the difference in ionization free energy between sequences where the *n*-neighbors can be either charged or uncharged according to :

$$\Delta\Delta G_{X,Z,n} = \frac{\Delta G_{Zz_{n-1}(X)z_{n-1}Z} - \Delta G_{z_n(X)z_n}}{2} \qquad (S18)$$

Where the uppercase Z is the charged state of the amino acid, and z is the uncharged state. We can then deduce the free energy for any context simply by adding the terms corresponding to each of the residues, as depicted in panel C in **Fig. 2**. For example, the ionization free energy of EA(K)KL:

$$\Delta G_{EA(K)KL} = \Delta G_{(K)} + \left(\Delta G_{expt} - \Delta G_{sim}\right) + \Delta\Delta G_{k,A,1} + \Delta\Delta G_{k,k,1} + \Delta\Delta G_{k,e,2} + \Delta\Delta G_{k,L,2} + \Delta\Delta G^{el}_{k,E,2} + \Delta\Delta G^{el}_{k,K,1} \qquad (S19)$$

Where $\Delta G_{expt} - \Delta G_{sim}$ is the difference in ionization free energy of the model compounds between our simulations and the values derived from experimental $pK_a$ measurments[2]. This term helps mitigate any systematic error due to the force field on our calculation. The parameters for window sizes 1 to 6 are shown in **Fig. S4** and **Fig. S5**. We note that Arg is not considered an ionizable amino acid, because of its $pK_a$ being extremely high (13.6[2,3]), and its high insensitivity to its environment[4,5]. As a result, Arg influence as a neighbor is always charged in our parameters. We demonstrate that this approach gives qualitatively similar results to the approach based on individual simulations of each separate context. However, the former approach greatly reduces the total number of simulations and allows the full database to be constructed before use on any specific protein, for much greater window sizes.

### Treatment of Histidine tautomers

For histidine, which has two tautomers in its neutral form, the free energy calculations from which the free energy of deprotonation are obtain for both the $H_\varepsilon$ and $H_\delta$ deprotonation. The overall free energy of deprotonation is obtained calculating the deprotonation free energy for each deprotonation tautomers, in a given context, and then combining them using equation 5. We note that to simplify the calculation, we treat the influence of neutral histidine on the free energy of deprotonation of another residue to be the same in both tautomers, and thus only use the dominant $H_\varepsilon$ as a context in the deprotonation free energy calculation.

**Time gains for the algorithm**

We can analyze the time gain from the use of each method by looking at the number of operations necessary for the full determination of the ensemble. In a typical evaluation, each of the $2^N$ microstates are generated and then evaluated. For each of these states, we need to add the relevant free energy of ionization for each of the residues that are in their deprotonated state. This cost is thus dependent on the specific microstates, but is constant within a mesostate. If we consider a mesostate $q$ with $n_q^{H^+}$ released protons and N ionizable residues, the number of operations to calculate the corresponding free energy of ionization addition $M_q$ is:

$$M_q = n_q^{H^+} \binom{N}{n_q^{H^+}} \qquad (S20)$$

This corresponds to obtaining the free energy of ionization for each released proton, for each microstate in this mesostate, the number of microstates for a given mesostate being the number of combinations of possible proton arrangements along the chain. Thus, the total number of operations is :

$$M_{tot} = \sum_{n_q^{H^+}=0}^{N} n_q^{H^+} \binom{N}{n_q^{H^+}} \qquad (S21)$$

This is depicted in by the red data in **Fig. 6**. If we simply use the iterative method discussed previously, the total number of operations becomes:

$$M_{tot} = \sum_{i=1}^{N+1} 2^i \qquad (S22)$$

That is, for each step of the procedure, we need keep track of $2^i$ residues states, until all ionizable residues have been accounted for. This is depicted by the blue data in **Fig. 6**. If we the dynamic ensemble building procedure introduced in this paper, the total number of operations is:

$$M_{tot} = n_{neigh}^2 \sum_{i}^{N} i \qquad (S23)$$

That is, we have $n_{neigh}^2$ contexts, each being added to all currently populated mesostates in the dynamic procedure. This is depicted by the green data in **Fig. 6**. If we choose to save the entire partition function on a per residue basis, then this equation becomes:

$$M_{tot} = N n_{neigh}^2 \sum_{i}^{N} i \qquad (S24)$$

This is depicted by the orange data in **Fig. 6**. This means that the total computational cost of determining the partition function is no longer scaling with $2^N$ but instead scales with $N^2$, which entails that the combinatorial explosion is no longer a problem when dealing with systems that have a large number of ionizable residues.

### Test of the maximum sequence length

We tested the maximum length of the sequence that could be tackled by MEDOC by using homopolymers of E. We found that the maximum number of Glu for which we could compute the full partition function was 925. This was done on an intel i7 2.2GHz, 16Gb of RAM machine running Ubuntu on python 3 64bits. We note that if not all software and dependencies are installed in a 64bit version, the limitation will be close to 130 residues, due to machine precision errors.

Details step by step description of the algorithm for EEE

**Step 1**

**1 q=0**

**0 q=-1**

$${}_{q=0}^{i=0}G_{s=1}^{p=1} = G_1 = \emptyset$$

$${}_{q=-1}^{i=0}G_{s=0}^{p=1} = G_0 = \Delta G_{\emptyset\emptyset(0)}$$

**Step 2**

| | | |
|---|---|---|
| **11** | | **q=0** |
| **10** | **01** | **q=-1** |
| **00** | | **q=-2** |

$${}_{all}^{i=1}G_{s=1}^{p=1} = f(G_{10}, G_{11}) = f\left(G_1 + \Delta G_{\emptyset 1(0)}, G_1 + \emptyset\right)$$

$${}_{q=0}^{i=1}G_{s=1}^{p=1} = G_{11}$$

$${}_{q=-1}^{i=1}G_{s=1}^{p=1} = G_{10}$$

$${}_{all}^{i=1}G_{s=1}^{p=2} = f(G_{01}, G_{11}) = f(G_0 + \emptyset, G_1 + \emptyset)$$

$${}_{q=0}^{i=1}G_{s=1}^{p=2} = G_{11}$$

$${}_{q=-1}^{i=1}G_{s=1}^{p=2} = G_{01}$$

$${}_{all}^{i=1}G_{s=0}^{p=1} = f(G_{01}, G_{00})$$

$${}_{q=0}^{i=1}G_{s=0}^{p=1} = +\infty$$

$${}_{q=-1}^{i=1}G_{s=0}^{p=1} = G_{01}$$

$${}_{q=-2}^{i=1}G_{s=0}^{p=1} = G_{00}$$

$${}_{all}^{i=1}G_{s=0}^{p=2} = f(G_{10}, G_{00})$$

$${}_{q=0}^{i=1}G_{s=0}^{p=2} = +\infty$$

$$ {}_{q=-1}^{i=1}G_{s=0}^{p=2} = G_{10} $$

$$ {}_{q=-2}^{i=1}G_{s=0}^{p=2} = G_{00} $$

**Step 3**

| | | | |
|---|---|---|---|
| **111** | | | **q=0** |
| **110** | **101** | **011** | **q=-1** |
| **100** | **010** | **001** | **q=-2** |
| **000** | | | **q=-3** |

$$ {}_{all}^{i=2}G_{s=1}^{p=1} = f(G_{110}, G_{101}, G_{100}, G_{111}) $$

$$ {}_{q=0}^{i=2}G_{s=1}^{p=1} = G_{111} $$

$$ {}_{q=-1}^{i=2}G_{s=1}^{p=1} = f(G_{110}, G_{101}) $$

$$ {}_{q=-2}^{i=2}G_{s=1}^{p=1} = G_{100} $$

$$ {}_{q=-3}^{i=2}G_{s=1}^{p=1} = +\infty $$

$$ {}_{all}^{i=2}G_{s=1}^{p=2} = f(G_{110}, G_{011}, G_{010}, G_{111}) $$

$$ {}_{all}^{i=2}G_{s=1}^{p=3} = f(G_{101}, G_{011}, G_{001}, G_{111}) $$

$$ {}_{all}^{i=2}G_{s=0}^{p=1} = f(G_{011}, G_{001}, G_{010}, G_{000}) $$

$$ {}_{q=0}^{i=2}G_{s=0}^{p=1} = +\infty $$

$$ {}_{q=-1}^{i=2}G_{s=0}^{p=1} = G_{011} $$

$$ {}_{q=-2}^{i=2}G_{s=0}^{p=1} = f(G_{001}, G_{010}) $$

$$ {}_{q=-3}^{i=2}G_{s=0}^{p=1} = G_{000} $$

$$ {}_{all}^{i=2}G_{s=0}^{p=2} = f(G_{101}, G_{100}, G_{001}, G_{000}) $$

$$ {}_{all}^{i=2}G_{s=0}^{p=3} = f(G_{110}, G_{100}, G_{010}, G_{000}) $$

### Implicit free energy rescaling

We used published site specific $pK_a$ values to compare our results. We originally found a qualitative agreement between the predicted and measured $pK_a$ which improved at high prediction temperatures, indicating an overestimation of the free energy differences due to the sequence context. This can be expected given the use of an implicit solvation force field for the obtention of these free energies, which can thus contain systematic biases and approximation. We thus chose to rescale the intrinsic and electrostatic contributions separately for each ionizable amino acids types, in order to improve agreement with experimental values. For this we use the relations :

$$\Delta\Delta G_{vol}^{rescaled} = \Delta\Delta G_{vol} \times S_{vol} + \text{offset} \qquad (S25)$$

$$\Delta\Delta G_{el}^{rescaled} = \Delta\Delta G_{el} \times S_{el} \qquad (S26)$$

Where $\Delta\Delta G_{vol}$ the is the intrinsic contribution to the free energy of deprotonation and $\Delta\Delta G_{el}$ is the electrostatic contribution, S is the corresponding rescaling factor. The value for the offset, and scaling factors are given in **Table S1** for the three residues that were included in the experimental dataset (E, D and H). The values for Y and K are computed as the average of the values for E, D and H. This scaling improved the agreement with experimental data, while conserving the amino acids specificity on the deprotonation free energy of both the protonatable residue and context residues. The resulting predicted vs experimental $pK_a$ values are shown in **Fig. 5C**.

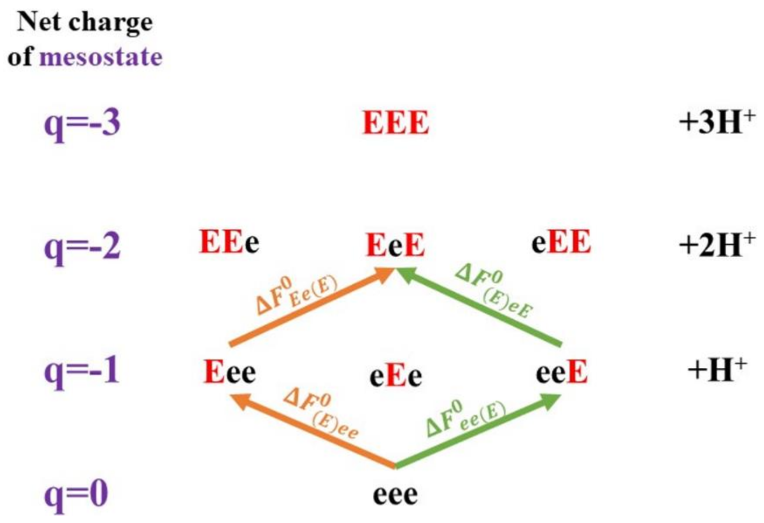


*Figure S1 : Schematic showing the thermodynamic path equivalence in between different ionization path for a eEe microstate from a Glu tripeptide.*

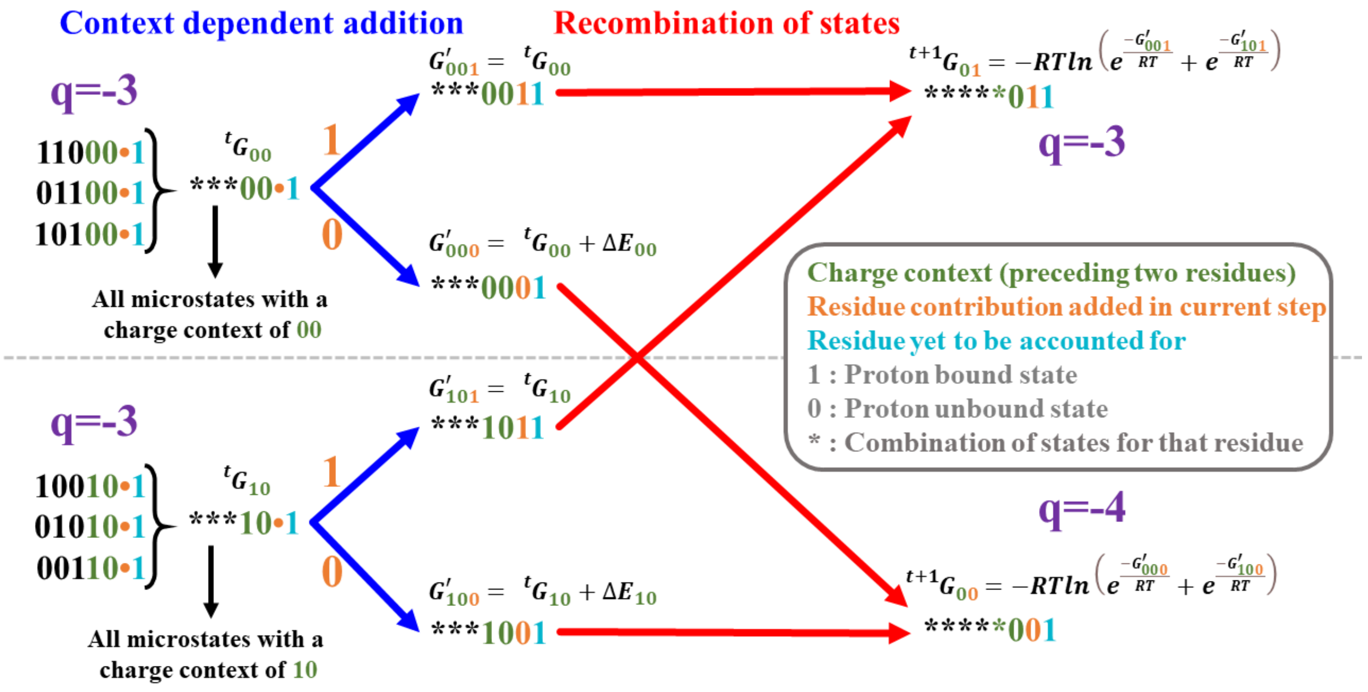


*Figure S2 : Schematic of the algorithm used to compute the ensemble weights without explicit representation of all microstates. 0 represents a residue having no bound proton, 1 represents a residue with a bound proton. A full description of the figure is available, in the S1 movie, which consists of the animated and voiced-over version of this figure.*

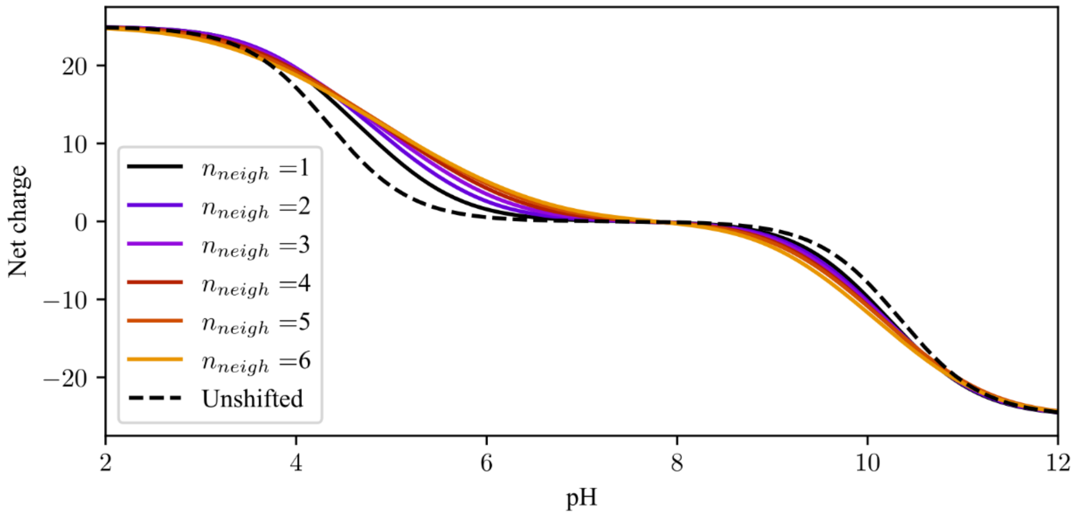


*Figure S3 : Effect of window size on the predicted net charge vs pH for a $E_{25}K_{25}$ amino acid sequence. The window size for each color is given in the legend*

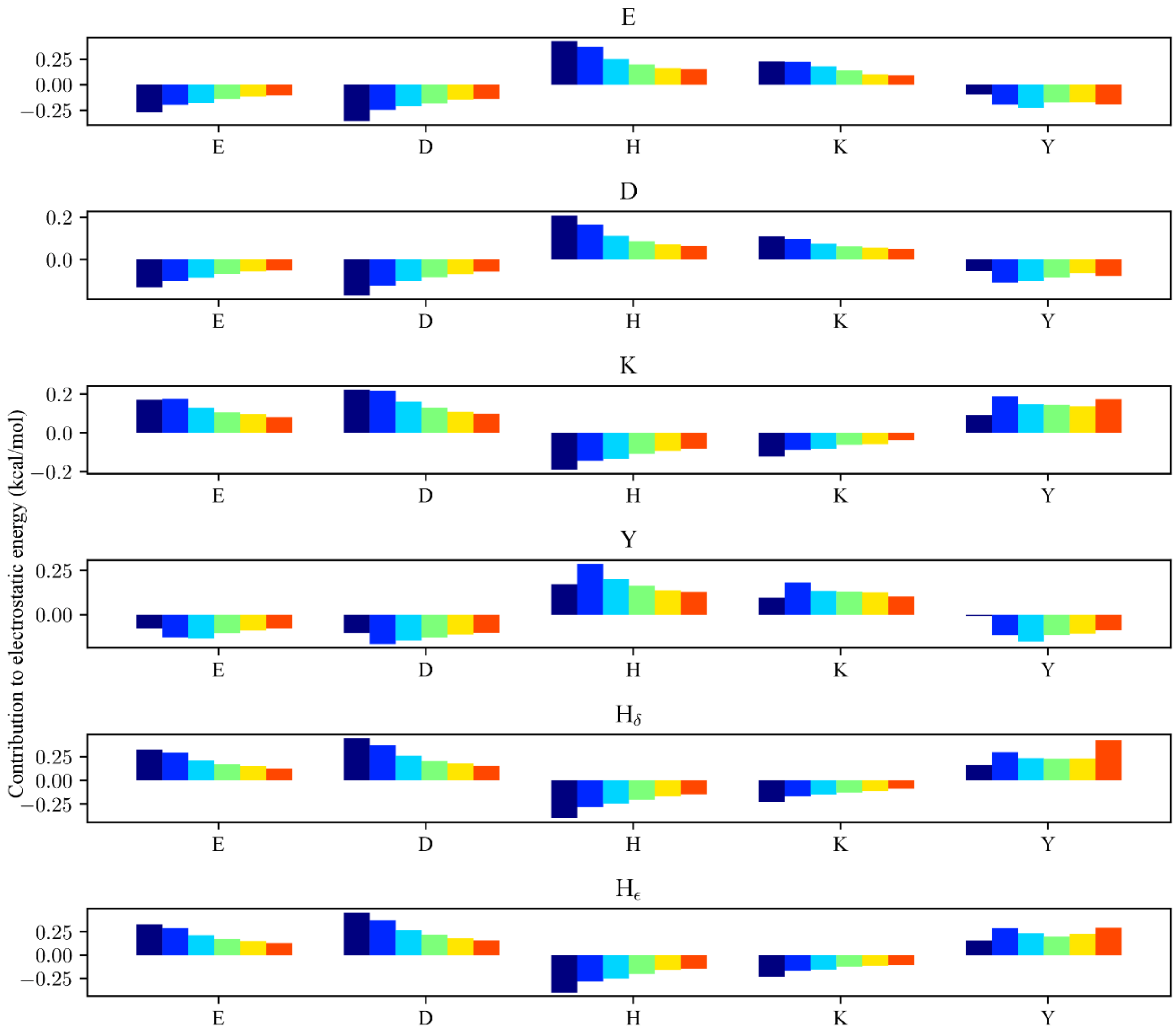


*Figure S4 : Implicit free energy determination electrostatic parameters. The residue indicated in the title of each graph is the residue being ionized, the residue indicated in the x axis for each graph is the neighbor residue. The parameters go from first neighbor (blue) to the 6th neighbor (orange).*

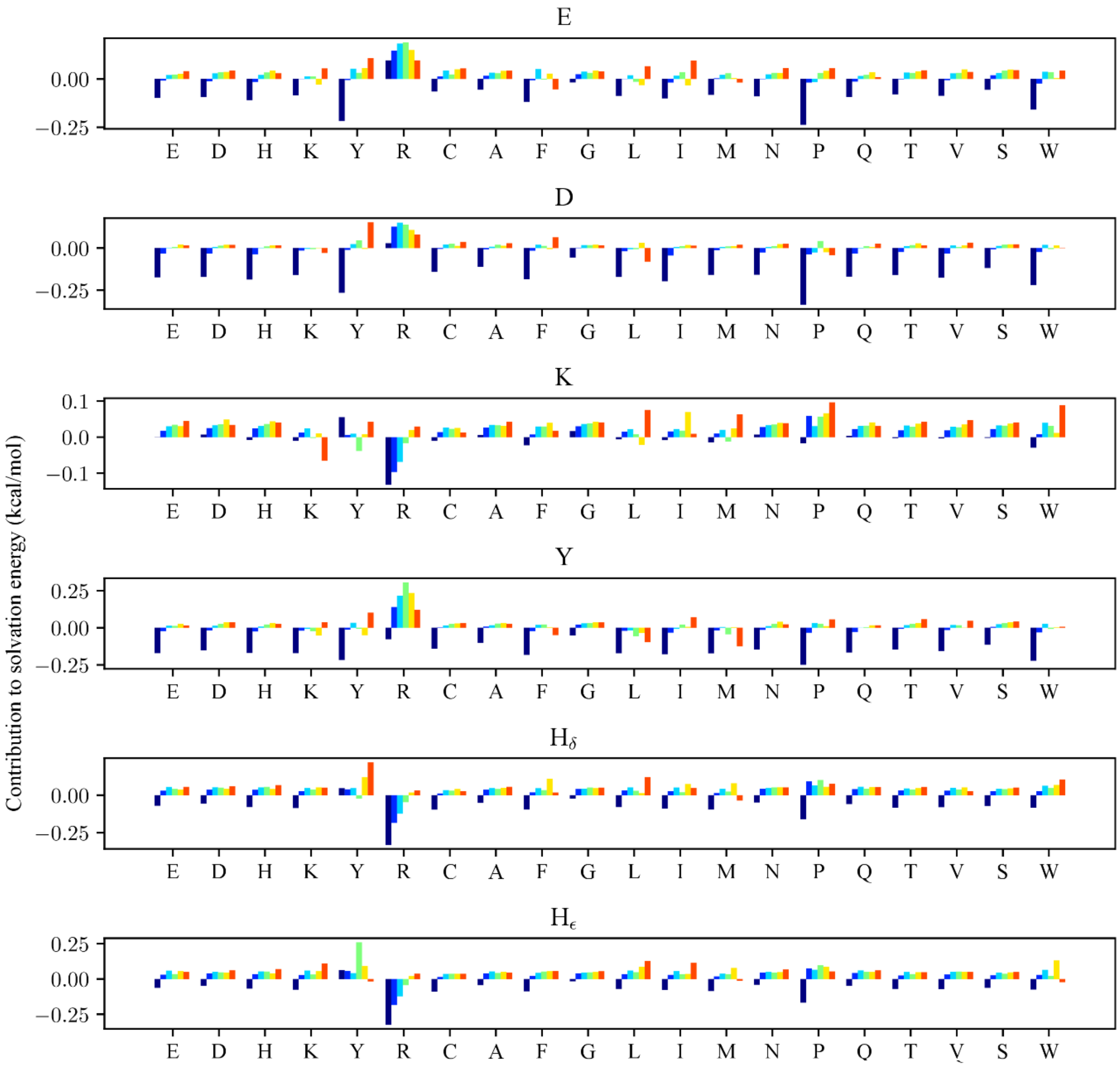


*Figure S5 : Implicit free energy determination solvation energy parameters. The residue indicated in the title of each graph is the residue being ionized, the residue indicated in the x axis for each graph is the neighbor residue. The parameters go from first neighbor (blue) to the 6th neighbor (orange). We note that since we do not consider R to be ionizable, this parameter contains the electrostatic energy within it.*

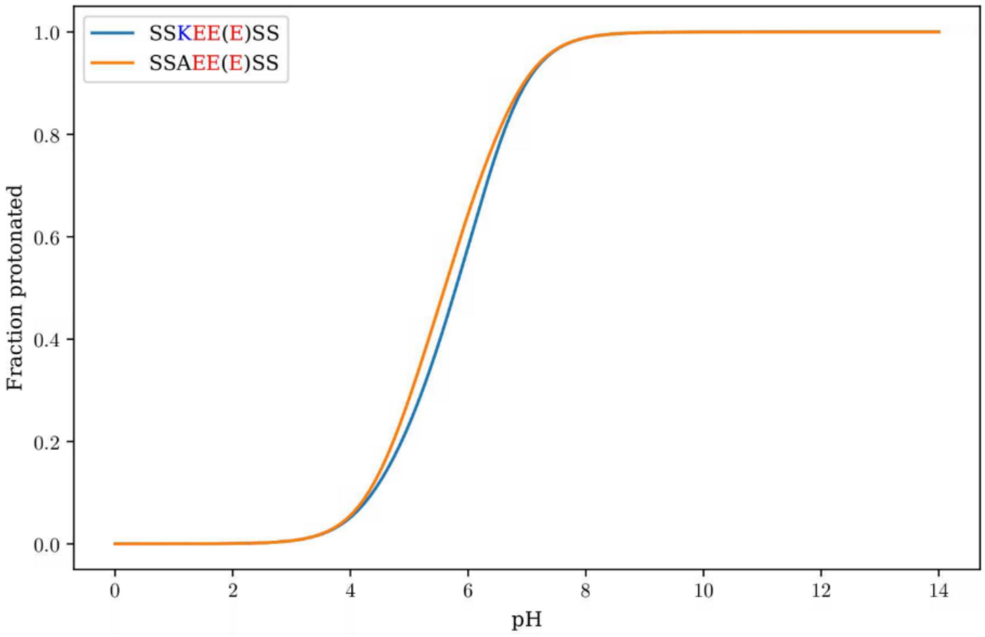


*Figure S6 : Depiction the effect of residues outside of the window on the fraction deprotonated of a residue, through a network of "connected" ionizable residues. Here the window size $n_{neigh}$=2. Each curve corresponds to the fraction deprotonated for the amino acid in parenthesis in the legend.*

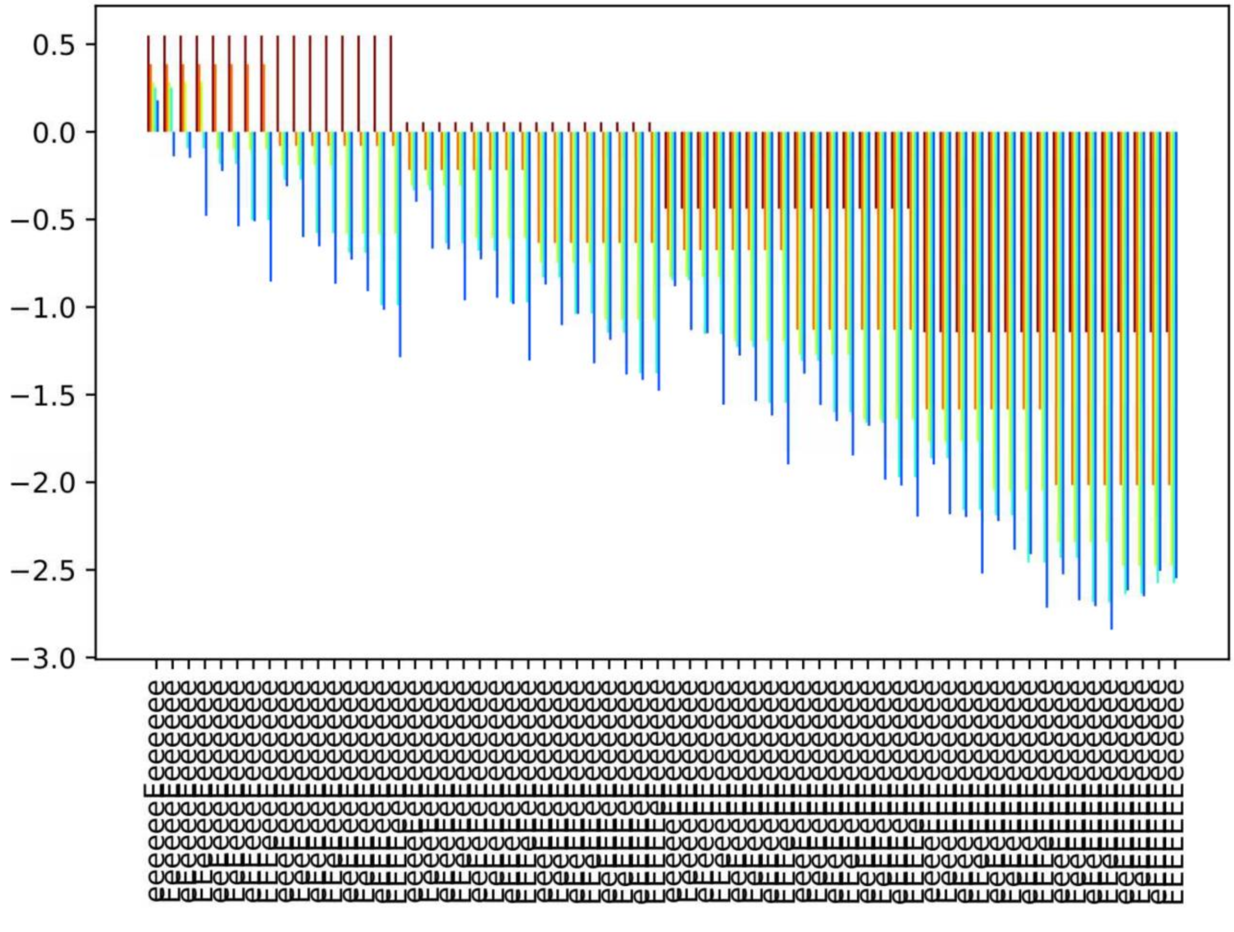


*Figure S7 : Effect of window size on microstates free energy. The window sizes go from 1 (maroon) to 6 (blue). The microstates here are show all 64 possible charge contexts of the 6 residues on the left (N-terminal) side of the amino acid for which the ionization free energy is assessed.*

*Table S1 : Rescaling factors and offset value for the deprotonation free energy rescaling procedure*

| | $S_{vol}$ | $Offset$ | $S_{el}$ scale |
|---|---|---|---|
| D | 0.30765 | 0.0221508 | 0.135366 |
| E | 0.408735 | 0.0461475 | 0.39555 |
| $H_{\varepsilon}$ and $H_{\delta}$ | 0.3809 | 0.054205 | 0.293 |
